# Turning high-throughput structural biology into predictive inhibitor design

**DOI:** 10.1101/2021.10.15.464568

**Authors:** Kadi L. Saar, Daren Fearon, The COVID Moonshot Consortium, Frank von Delft, John D. Chodera, Alpha A. Lee

## Abstract

A common challenge in drug design pertains to finding chemical modifications to a ligand that increases its affinity to the target protein. An underutilised advance is the increase in structural biology throughput, which has progressed from an artisanal endeavour to a monthly throughput of up to 100 different ligands against a protein in modern synchrotrons. However, the missing piece is a framework that turns high throughput crystallography data into predictive models for ligand design. Here we designed a simple machine learning approach that predicts protein-ligand affinity from experimental structures of diverse ligands against a single protein paired with biochemical measurements. Our key insight is using physics-based energy descriptors to represent protein-ligand complexes, and a learning-to-rank approach that infers the relevant differences between binding modes. We ran a high throughput crystallography campaign against the SARS-CoV-2 Main Protease (M^Pro^), obtaining parallel measurements of over 200 protein-ligand complexes and the binding activity. This allows us to design a one-step library syntheses which improved the potency of two distinct micromolar hits by over 10-fold, arriving at a non-covalent and non-peptidomimetic inhibitor with 120 nM antiviral efficacy. Crucially, our approach successfully extends ligands to unexplored regions of the binding pocket, executing large and fruitful moves in chemical space with simple chemistry.

## Introduction

Predicting protein-ligand affinity is a longstanding challenge that underpins computer-aided drug design. The challenge often lies in designing chemical modifications which would significantly improve the potency of a weakly potent starting point (hit-to-lead) or finding chemotypes that maintains potency whilst designing away other liabilities (lead optimisation). Established medicinal chemistry heuristics focus on making interpretable and modest chemical changes, iteratively “morphing” the ligand to optimise interactions and explore unknown binding pockets ^1,2^. Significant acceleration can be realised if this iterative process is replaced by methods which suggest large and synthetically facile changes to the ligand to give a significant increase in potency, motivating a computational approach to ligand design.

The plethora of computational methods in the literature can be organised in terms of the available information they make use of. Ligand-based approaches (Figure a) derive information only from the chemical identity of ligands which are binding to the protein, and focus on learning the relationship between the chemical structure of the ligand and its activity. Such methods, however, are circumscribed by the problem of extrapolation: the model cannot extrapolate to regions of the binding site which are not already explored by molecules in the dataset, nor to unexplored interactions between novel chemotypes and the binding site.

Structure-based approaches ameliorate this limitation by taking the protein structure into account and explicitly model the protein-ligand interactions (Figure b). However, rigorous methods, such as free energy perturbations (FEP) or alchemical free energy calculations generally require substantial computational resources and are constrained by the quality of the approximate forcefield. ^3–5^ In practice, these approaches are typically used to compute relative free energy changes of small modifications to predefined scaffolds ^6,7^, as convergence time and error both increase with the size of change relative to the starting ligand. To reduce computational cost, empirical scoring functions such as docking ^8–11^ have been developed. Whilst docking can identify hits from virtually screening large libraries ^12,13^, it is typically not used in ligand optimisation as the correlation between predicted and experimentally measured protein-ligand interaction energy is often weak.

**Figure 1:**
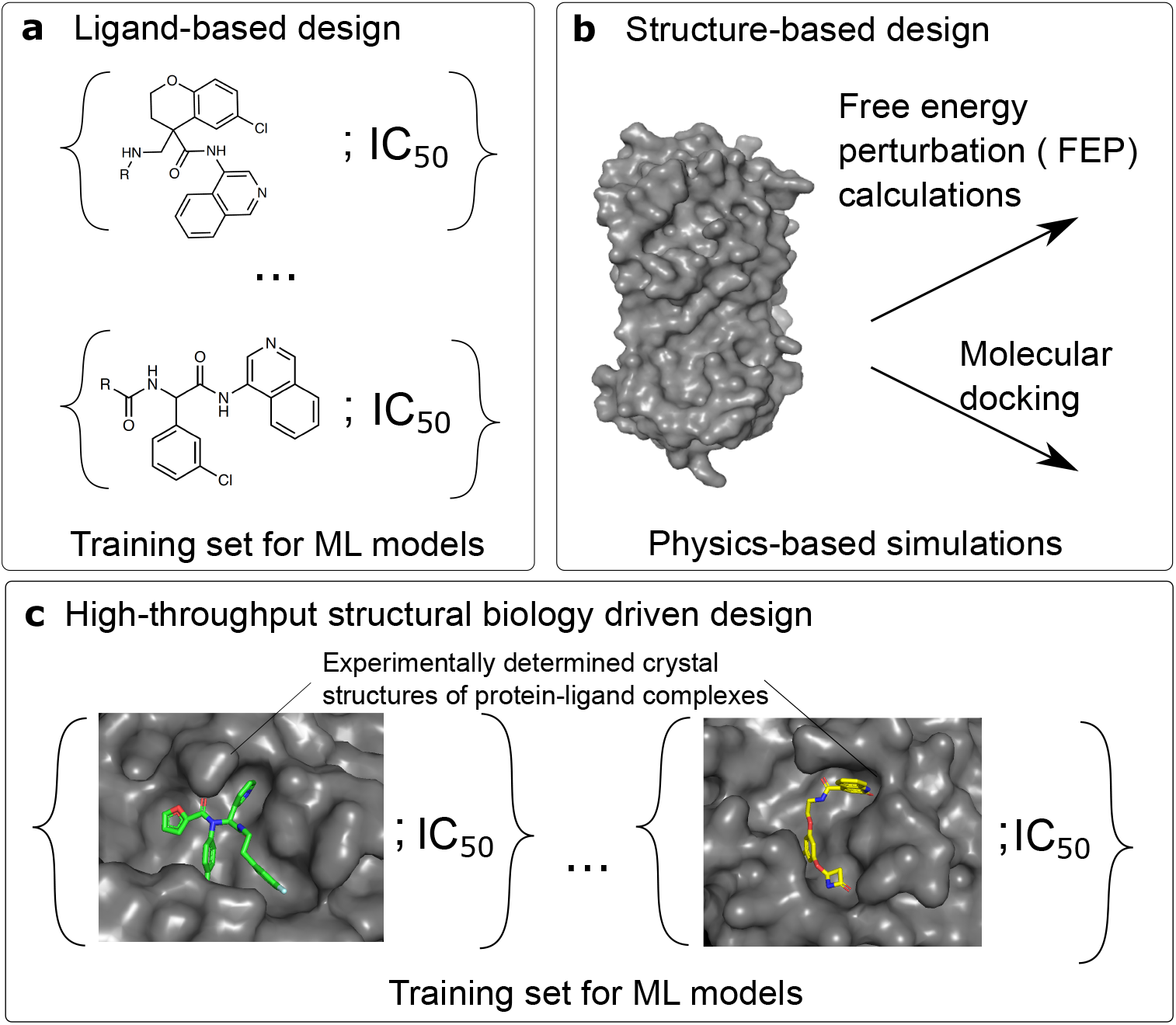
Overview of approaches used in computer-aided drug design. Conventional approaches predict the activity of ligands **(a)** by learning the relationships between the molecular structure of a molecule and its activity or **(b)** through physics-based modelling using only the structure of the target and relying on methods such as free energy perturbation (FEP) calculations or molecular docking. **(c)** Here, we demonstrated a strategy that exploits high-throughput crystallographic characterisation of protein-ligand complexes to predict ligand affinity.

The rapid acceleration in the throughput of structural biology unlocks a new source of data (Figure c). Historically, protein structure determination was laborious, thus on a particular target there were only handful of co-crystallised ligands reported in the literature. Although databases such as the Protein Data Bank ^14^ could be mined to parametrize docking algorithms ^15,16^, this necessitates training on diverse classes of proteins with varied protein-ligand affinity measurement techniques, introducing noise and dataset bias ^17^. The synergy between modern robotic techniques for crystallisation and crystal soaking ^18^, automated data analysis pipelines ^19^ and modern synchrotron infrastructure has increased the monthly throughput to up to 100s of ligands against a target ^20^. However, the missing piece of the puzzle is a framework that can turn high throughput crystallographic data into predictive models for ligand design.

In this paper, we present a machine learning approach that relates high throughput crystallography data, represented as empirical energy terms, to measured bioactivity. We used this to accelerate the COVID Moonshot initiative ^21^, an open science consortium that reported over 200 protein-ligand complexes against SARS-CoV-2 Main Protease (M^Pro^) with associated potency (*IC*_50_) measurements. Retro-spective validation shows that our method outperforms ligand-based and structure-based approaches. We prospectively designed one-step library syntheses, improving the potency of two distinct micromolar hits by over 10-fold and arrived at a lead compound with 120 nM antiviral efficacy. Crucially, our designed inhibitors gain potency by extending to unsampled regions of the binding site, illustrating the ability of our model to generalise via the incorporating physical interactions.

## Results and Discussion

### Energy-based model is generalizable across chemical space

To describe protein-ligand structures as a fixed-length vector for downstream machine learning (Figure a), we turn to the literature on empirical scoring functions. We use the terms of empirical energy function – hydrophobic, electrostatic, hydrogen bonding etc. – as descriptors of the structures. Our hypothesis is that whilst empirical energy terms capture different aspects of protein-ligand interactions, how these interactions stack up to yield the free energy of binding depends on binding-site specific variables such as binding site flexibility. We further hypothesize that those protein-specific corrections are learnable from our dataset, comprising high throughput structural biology data and associated bioactivity. To fix ideas, in our approach we featurize all the protein-ligand complexes using the Open Drug Discovery Toolkit (ODDT) ^22^ and extract the Autodock Vina descriptors to serve as a low-dimensional representation of the each structure.

We predict protein-ligand affinity using this representation as an input. Instead of predicting IC_50_ values directly, we focused our attention to predicting the pairwise comparison of the ligands, choosing a cut-off of 0.25 log_10_ units for classifying one compound as more active than another (Figure S1). The threshold was chosen to match typical assay error. This learning-to-rank approach allows us to combine qualitative (potency below measurable) and quantitative measurements, and forces the model to ignore irrelevant experimental noise by ensuring that it is only ranking structures with demonstrably different bioactivity ^23^. We build our models using Autodock Vina descriptors as features and a random forest as the machine learning algorithm (Materials and Methods; Supplementary Figure S1).

Specifically, we applied this approach to the high-throughput structural biology campaign against the SARS-CoV-2 M^Pro^, an essential protein in viral replication and a validated target for anti-coronavirus therapeutics ^24–27^. All M^Pro^ clinical candidates to date are peptidomimetics inhibiting via a covalent mechanism, which are generally suboptimal for drug development. We launched the COVID Moonshot, an open science initiative aiming to develop non-covalent small molecule oral antiviral ^21^. The campaign obtained 236 structures of non-covalent inhibitors binding to the M^Pro^. Out of these ligands, 94 had IC_50_ below 50 *μ*M (Figure S2). To the best of our knowledge, COVID Moonshot is the only openly accessible dataset with over 100 structures of different ligands against a single target with associated bioactivity measurement; as such, our model evaluation will focus on this dataset.

To implement the models and evaluate their performance in ranking novel ligands, first, we use a scaffold-split approach where an entire scaffold is held out from the training set and placed in the test set. There are four salient chemical scaffolds in the dataset: aminopyridine-like, isoquinoline, benzotriazole and quinolone (Materials and Methods; Supplementary Figure S2) with 123, 44, 19 and 15 structures, respectively. For the compounds that were part of the test set, we extracted the features from the docked structures instead of the experimental crystal structures, where the ligands had been docked to the active site using OpenEye’s FRED hybrid docking mode as implemented in the “Classic OEDocking” floe on the Orion online platform (Materials and Methods). This is because when deploying the model as a prioritisation tool for synthesis and screening as we will do later in this work, experimental structural information is inaccessible and the docked structure serve as its approximation. Figure b shows that our learn-to-rank model achieved an average AUROC value around 0.8 (Supplementary Figure S3 shows the results broken down into different scaffolds). To estimate error in the performance metric, the training data was bootstrapped and 10 different models built for each scaffold, the quoted values are the mean and the standard deviations in the obtained AUROC values. This result illustrates that our model accurately ranks unseen ligands without the requirement to have any structures from that specific scaffold as part of the training set.

**Figure 2:**
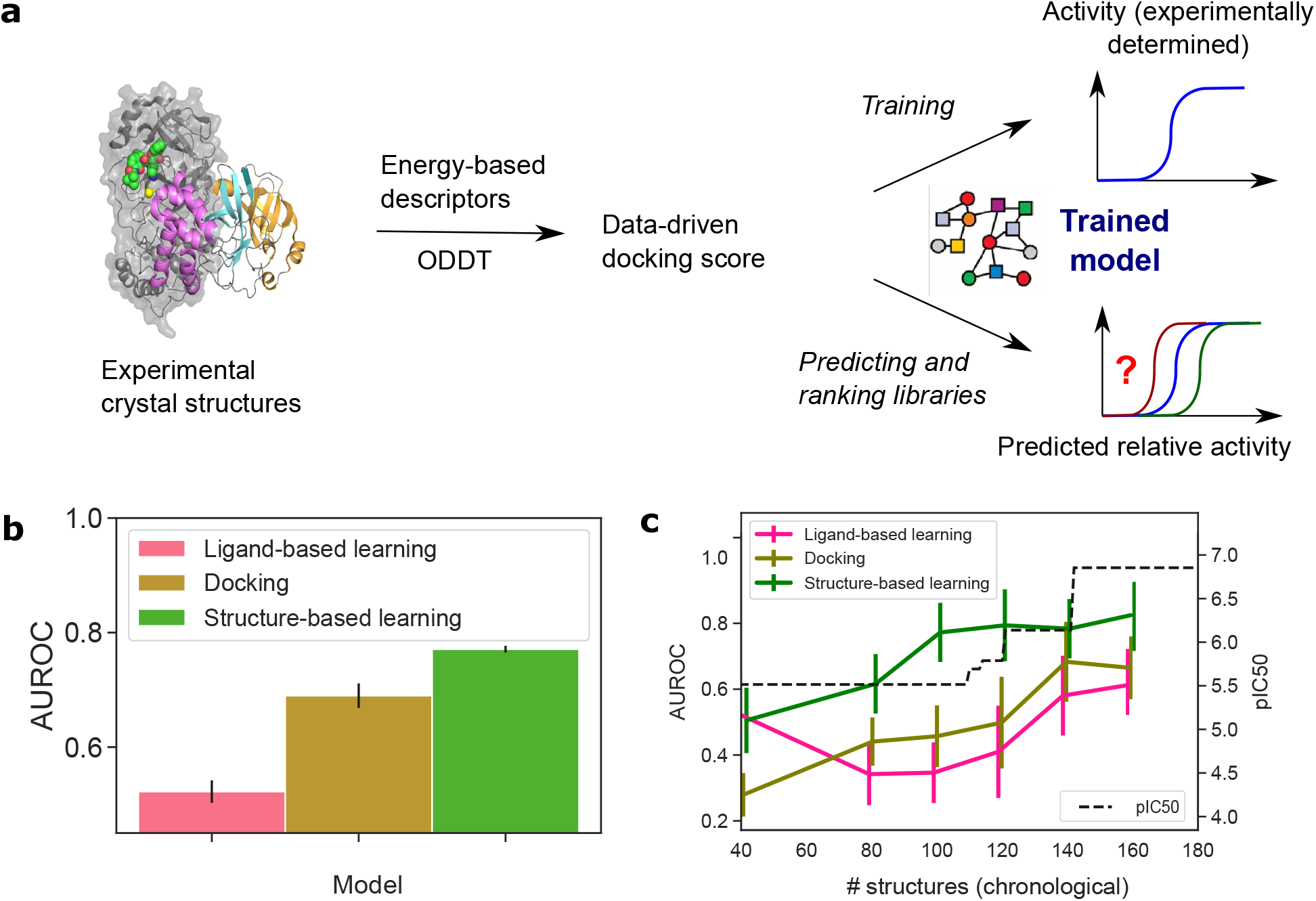
Structure-based learning outperforms docking and ligand-based machine learning in relative affinity predictions. **(a)** To build a model that captures the relationship between the crystal structure of a protein-ligand complex and the activity of the ligand, Autodock Vina descriptors were extracted from each crystal structure and deployed to predict the relative binding strength of ligands of interest. **(b)** Comparing this approach (green) to ligand-based learning (pink) and docking (yellow) using data from the COVID Moonshot campaign. The average AUROC scores were computed using a scaffold-split, with the error bars corresponding to standard error of the mean. Performance on individual scaffolds is shown in Supplementary Figure S3. **(c)** Time-split evaluation of our model. Our approach maintains its predictive power when trained on older, less potent molecules and asked to rank the affinities of newer more potent molecules (green line). The black line tracks the potency of most potent molecule discovered at that point.

### Structural data is salient to model performance

To understand the impact of experimental structural biology data, we consider two alternative models: (i) a ligand-based model that relied only on the use of ligand-based descriptors providing no information about the protein-crystal structure and (ii) a model that used a docked structure instead of the measured crystallographic structure. Specifically, for the former case, we featurised the ligands using Morgan fingerprints ^28^, implemented through the use of the RDKit package. For the latter case, we docked ligands to the active site using OpenEye’s FRED hybrid docking modes described above. For consistency, the same model architecture (random forest) was used for all cases with the hyperparameters tuned separately for each model.

Figure b shows that our approach which incorporates experimental structural data outperforms both docking-based and ligand-based models, highlighting the importance of high throughput structural biology. The AUROC values correspond to the average values across the four scaffolds. The slightly better performance of the docking-based model over the ligand-based one likely stems from the fact that the model does not rely solely on ligand-based input but also incorporates information about the protein-ligand interactions.

### Model maintains performance throughout the campaign

Having demonstrated that our proposed strategy can reliably rank ligands by potency even when outside the chemical space that it encountered during training (i.e. for a new scaffold), we next explore how its utility varies through the campaign and what is the amount of structural data required for efficient performance. As a drug discovery campaign progresses, more knowledge about the chemical attributes that determine the binding of a ligand to its target is gathered and the designs are honed accordingly. Therefore, trying to predict the potency of molecules tested earlier in the campaign with molecule tested later in the campaign as training set is much easier (and less useful) than the converse, i.e. hindsight is usually much more accurate than foresight.

To examine the predictive capability of the structure-driven ligand prioritisation approach as the campaign progresses, we use a time-split strategy. We ordered the compounds by the time when they had been tested and used only the structures available until that specific time point for training. We first note that in the course of the campaign, the potency of the molecules increased by many orders of magnitude (Figure c, black line). To avoid susceptibility of the model to memorise the specifics of a particular scaffold, we kept all aminopyridine-like and isoquinoline-like molecules in the training data while using benzotriazole-like and quinolone-like ligands for validation. Figure c (green line) shows that our structure-based model remains predictive when trained on molecules tested early in the campaign and deployed to rank molecules tested later. This is in contrast to the model that relied on ligand-based or docking-based input (Figure c, pink and dark yellow lines).

Finally, from these data, we can interrogate that our proposed approach performs effectively (AU-ROC value above 0.7) when only around 100 crystal structures are available to train the model. Crucially, this is a throughput that could be achieved in a modern synchrotron ^20^ on monthly timescale, illustrating that our proposed strategy has the potential to be exploited routinely in the context of drug discovery campaigns.

### Model-guided library synthesis discovers potent leads

To apply our model to lead discovery, we need to generate protein-ligand structures for unseen ligands. Starting from two hits with a amine handle reported by the COVID Moonshot Consortium ^29^, chosen because they have detectable potency and ease of synthetic access, we generate a virtual library that is synthesizable in single reaction step using amide formation (Figure 3a) and reductive amination (Figure 3b). The library design is motivated by structural data, aiming to extend the hit into the unoccupied P1’ binding pocket (Figure 3, top left). Using the Manifold platform (postera.ai/manifold), we select carboxylic acid and aldehydes that are in-stock building blocks in Enamine (a synthetic chemistry CRO with one of the largest building block collections onsite) with the building blocks further filter based on predicted reactivity and the final compound having *c*log*P* < 3. In total, there are 15,720 compounds in the amide virtual library and 2,664 compounds in the reductive amination library.

**Figure 3:**
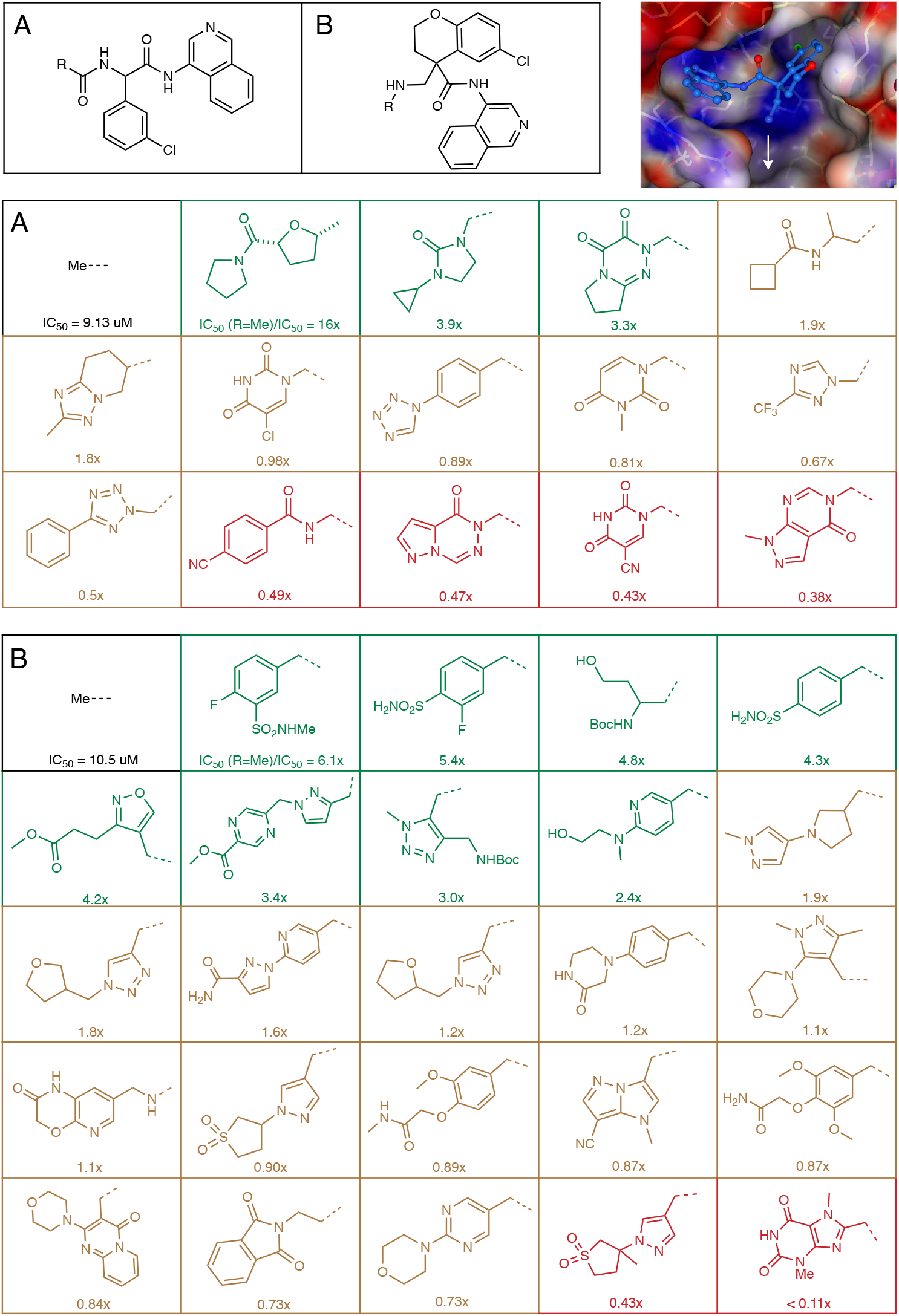
Lead compounds are starting points for antiviral development. Top hits reported in Figure 3 is purified and assayed in enzymatic assay. Compound 1, with nM potency, is further profiled in SARS-CoV-2 CPE assay in Vero E6 cells. The 95% CI for the enzymatic assays: Compound 1 [0.23, 0.43] *μM*, Compound 2 [0.96, 1.2] *μM*, Compound 3 [1.3, 1.7] *μM*.

We then generated predicted binding pose by constrained docking into the binding site using existing structural data in the isoquinoline series as the constraints (Materials and Methods), and use our trained structure-based learn-to-rank approach to rank all the ligands in the virtual library. Specifically, each of the docked poses are ranked against the top 5 most potent non-covalent binders in the dataset (Supplementary Figure S4) and the mean of the five predictions estimated to generate the final ranking. Top 18 compounds from the final ranking were selected from the amide formation library with 15 successfully synthesized, and top 32 from reductive amination library with 24 successfully synthesized.

The compounds prioritised by our crystallography-driven approach were then assessed for Inhibition of M^*pro*^ activity using a biochemical assay with a fluorescence-based readout ^23,29^. Figure 3 shows that around 30% of the library has potency that is that greater than 2x compared to the reference. These compounds are all substantial changes to the hit, in some cases doubling the atom count, and reaching the ligand into unknown regions of the binding site whilst remaining a low log*P*. The high hit rate suggests that the model can accurately prioritise these new chemotypes. We note that compounds were assayed as enantiomers or diastereoisomers in this initial triage.

We further characterise the potent leads by resolving the enantiomers/diasteroisomers. Figure 4 shows that our top enantiopure compound, Compound **1**, achieves nM potency in florescence Mpro assay. Compound **1** is further profiled in SARS-CoV-2 antiviral assay (CPE assay, in Vero E6 cells), attaining *EC*_50_ = 120nM. As a control, Compound **1** displays no cytotoxicity effect against Vero E6 cells at 10*μ*M. Compound **1** is thus a starting point for the development of antiviral therapeutics.

**Figure 4:**
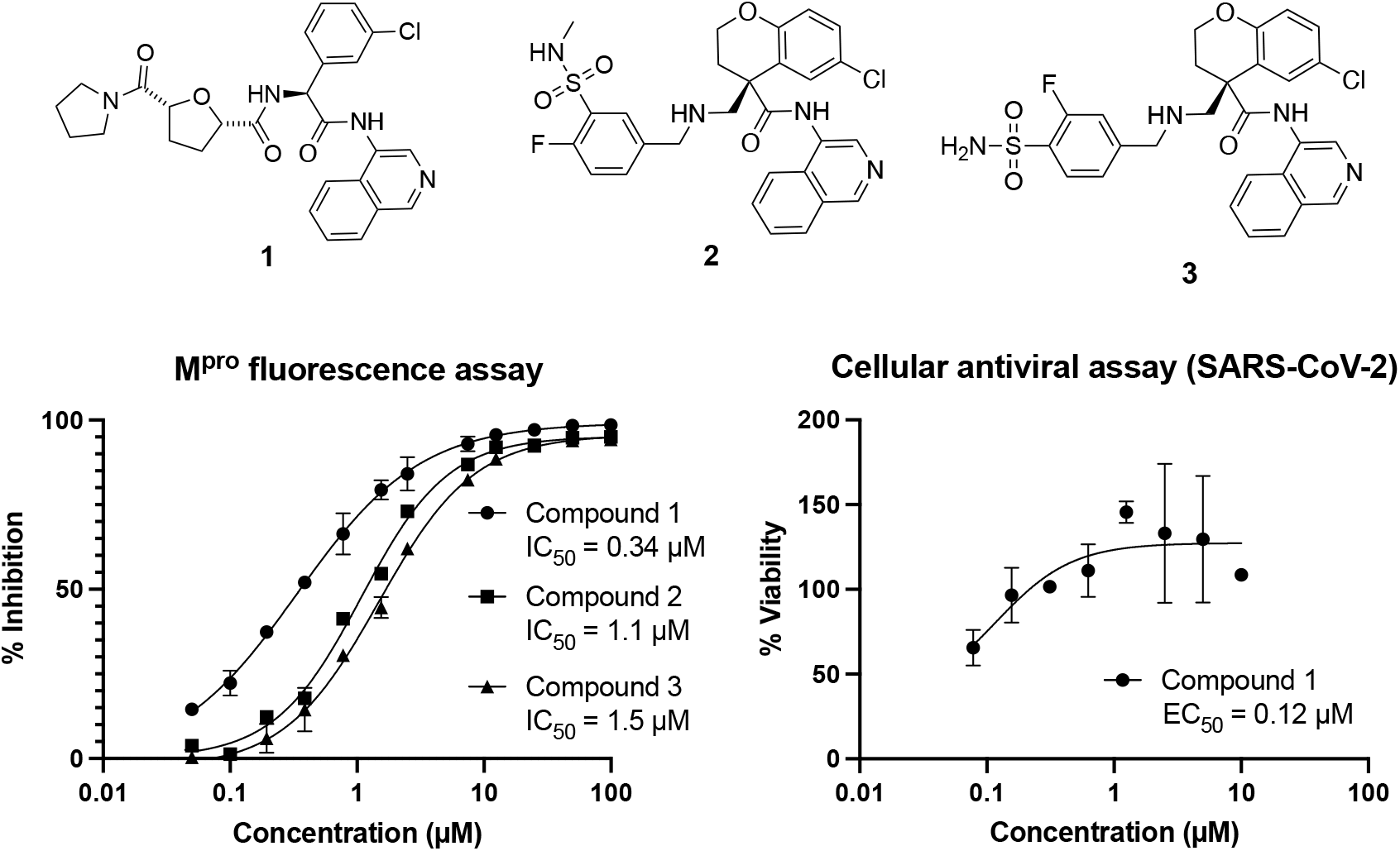
Lead compounds are starting points for antiviral development. Top hits reported in Figure 3 are purified and assayed in enzymatic assay for their activity against the SARS-CoV-2 main protease. Compound 1, with nM potency, is further profiled in SARS-CoV-2 CPE assay in Vero E6 cells and found to show no cytotoxic effect at 10 *μ*M. The 95% CI for the enzymatic assays: Compound 1 [0.23, 0.43] *μ*M, Compound 2 [0.96, 1.2] *μ*M, Compound 3 [1.3, 1.7] *μ*M.

## Conclusion

With the crystal structures of protein-ligand complexes being acquired at an increasing throughput, here we show how these data can be used to power a new approach to computational ligand design by using physics-inspired empirical energy terms as a descriptor of the protein-ligand complex. We focus on the COVID Moonshot Initiative, which reported an unpredentedly rich dataset of 200 ligands for which both their activity and structure of binding to the main protease of SARS-CoV-2 had been determined. We developed a machine learning model that learned the relationship between the multi-dimensional docking score extracted from the crystal structure and the relative bioactivity of ligands. The approach maintained a high and robust performance (AUROC of 0.79), even when making predictions outside the training scaffold. It also yielded powerful results in a prospective campaign, increasing the potency of hit compounds by more than 10x with simple chemistry that extends the hits to unsampled region of the binding site. Our approach arrived at a lead compound with 120 nM antiviral efficacy.

## Materials and Methods

### Code availability

Source code and data required to reproduced this study are available on: https://github.com/kadiliissaar/ligand_design_structural_biology

### Dividing the molecules by scaffolds

In order to reliably estimate the performance of our developed model, we performed the train:test splits in a scaffold stratified manner. To this effect, four distinct scaffold categories were defined — aminopyridine-like, isoquinoline, benzotriazole and quinolone — and each compound classified as belonging into one of them by using SMARTS to define chemical substructures. Figure S5 shows representative examples of each chemical series, highlighting with the chemical substructure that gives rise to the name.

### Model development

As described in the Main Text, two types of models were developed: (i) those that divided the full dataset into the four scaffolds and developed four parallel models by each time keeping molecules that originated from one of the four scaffolds as a test set and using the remainder of the data for training, and (ii) those that sorted all the data chronologically by the date the molecule was tested.

In all cases, after splitting the data into the training and test sets, all possible pairs were generated and a difference in the pIC_50_ value as well as between the descriptors evaluated as as been illustrated in Supplementary Figure S1. Compounds that were determined to be inactive were included when forming pairs and they were all allocated activities equal to the highest measured activity values. Following this data preparation step, a random forest classifier was trained to build a model that would predict if the difference in the pIC_50_ value is above (the activity of the ligands differs) or below (the activity of the ligands does not differ) 0.25 log_10_ units using the difference in the descriptors extracted from the structures of the respective compounds as the input features. Hyperparameters of each model were tuned by performing a 10-fold cross-validation process and a random grid search. Generally, the best performance was achieved at shallow forest depths (maximum depth of 3), which can be explained by the relatively large number of training points (order of 10^3^ - 10^4^) in comparison to the number of features (6).

### Docking experiments

We redocked all compounds synthesized by The COVID Moonshot Consoritium against x2908 structure reported by Diamond XChem. We use the “Classic OEDocking” floe v0.7.2 as implemented in the Orion 2020.3.1 Academic Stack (OpenEye Scientific). Omega was used to enumerate conformations (and expand stereochemistry) with up to 500 conformations. FRED was used for docking in HYBRID mode using the x2908 bound ligand. The docked poses are available on GitHub.

### Fluorescence MPro inhibition assay

Method is described previously ^29^. Compounds were seeded into assay-ready plates (Greiner 384 low volume, cat 784900) using an Echo 555 acoustic dispenser, and DMSO was back-filled for a uniform concentration in assay plates (DMSO concentration maximum 1%) Screening assays were performed in duplicate at 20*μM* and 50*μM*. Hits of greater than 50% inhibition at 50*μM* were confirmed by dose response assays. Dose response assays were performed in 12 point dilutions of 2-fold, typically beginning at 100*μM*. Highly active compounds were repeated in a similar fashion at lower concentrations beginning at 10*μM* or 1*μM*. Reagents for Mpro assay were dispensed into the assay plate in 10μl volumes for a final volume of 20*μM*.

Final reaction concentrations were 20mM HEPES pH7.3, 1.0mM TCEP, 50mM NaCl, 0.01% Tween20, 10% glycerol, 5nM Mpro, 375nM fluorogenic peptide substrate ([5-FAM]-AVLQSGFR-[Lys(Dabcyl)]K-amide). Mpro was pre-incubated for 15 minutes at room temperature with compound before addition of substrate and a further 30 minute incubation. Protease reaction was measured in a BMG Pherastar FS with a 480/520 ex/em filter set. Raw data was mapped and normalized to high (Protease with DMSO) and low (No Protease) controls using Genedata Screener software. Normalized data was then uploaded to CDD Vault (Collaborative Drug Discovery). Dose response curves were generated for IC50 using nonlinear regression with the Levenberg–Marquardt algorithm with minimum inhibition = 0% and maximum inhibition = 100%.

### SARS-CoV-2 antiviral assay

Method is described previously ^29^. SARS-CoV-2 (GISAID accession EPI ISL 406862) was kindly provided by Bundeswehr Institute of Microbiology, Munich, Germany. Virus stocks were propagated (4 passages) and tittered on Vero E6 cells. Handling and working with SARS-CoV-2 virus was conducted in a BSL3 facility in accordance with the biosafety guidelines of the Israel Institute for Biological Research (IIBR). Vero E6 were plated in 96-well plates and treated with compounds in medium containing 2 % fetal bovine serum. The assay plates containing compound dilutions and cells were incubated for 1 hour at 37oC temperature prior to adding Multiplicity of infection (MOI) 0.01 of viruses. Viruses were added to the entire plate, including virus control wells that did not contain test compound and Remdesivir drug used as positive control. After 72h incubation viral cytopathic effect (CPE) inhibition assay was measured with XTT reagent. Three replicate plates were used.

### Chemical Synthesis

All compounds were synthesized at Enamine and available for purchase from their catalogue. Detailed description of the synthesis protocol is outlined in the Supplementary Materials.

## Supporting information

Supplementary Materials - Chemical Synthesis

## Acknowledgements

The research leading to these results has received funding from the Schmidt Science Fellows program in partnership with the Rhodes Trust (KLS). The Chodera laboratory receives or has received funding from multiple sources, including the National Institutes of Health, the National Science Foundation, the Parker Institute for Cancer Immunotherapy, Relay Therapeutics, Entasis Therapeutics, Silicon Therapeutics, EMD Serono (Merck KGaA), AstraZeneca, Vir Biotechnology, Bayer, XtalPi, Interline Therapeutics, the Molecular Sciences Software Institute, the Starr Cancer Consortium, the Open Force Field Consortium, Cycle for Survival, a Louis V. Gerstner Young Investigator Award, and the Sloan Kettering Institute. A complete funding history for the Chodera lab can be found at http://choderalab.org/funding. AAL holds a Royal Society University Research Fellowship.

## Author contributions

KLS and AAL designed the study. DF, FvD and the COVID Moonshot Consortium performed all structural characterisation work, biochemical and antiviral assays. JCD designed and performed the docking experiments. KLS and AAL devised the predictive model and KLS implemented it. KLS and ALL wrote the original draft, all authors commented on it.

