## Supplementary Materials - Chemical Synthesis for "Turning high-throughput structural biology into predictive inhibitor design"

**The Synthesis of rel-(2S,5R)-N-((R)-1-(3-chlorophenyl)-2-(isoquinolin-4-ylamino)-2-oxoethyl)-5-(pyrrolidine-1-carbonyl)tetrahydrofuran-2-carboxamide (Z5056159151, H3319657):**

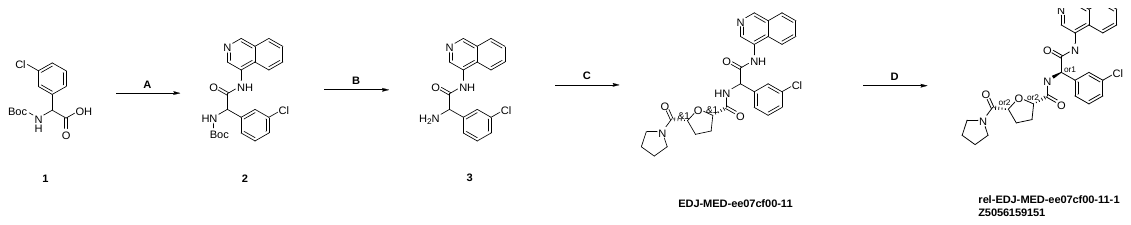

**Step A:**

CC(C)(C)OC(=O)NC(C(=O)O)C=1C=CC=C(Cl)C1.(Cl.NC=1C=NC=C2C=CC=CC12).CN(C)C=1C=CN=CC1.(Cl.CCN=C=NCCCN(C)C).CCN(C(C)C)C(C)C>>CC(C)(C)OC(=O)NC(C(=O)NC=1C=NC=C2C=CC=CC12)C=3C=CC=C(Cl)C3

2-[(tert-Butoxy)carbonyl]amino-2-(3-chlorophenyl)acetic acid (7.0 g, 24.48 mmol), isoquinolin-4-amine hydrochloride (4.02 g, 22.26 mmol), N,N-dimethylpyridin-4-amine (543.86 mg, 4.45 mmol), N-(3-dimethylaminopropyl)-N'-ethylcarbodiimide hydrochloride (6.4 g, 33.39 mmol), ethylbis(propan-2-yl)amine (6.33 g, 48.97 mmol, 8.53 ml, 2.2 equiv) were suspended in DMF and heated at 50 ºC overnight. After cooling to room temperature, the mixture was evaporated, and the residue was mixed with water. The obtained mixture was filtered and the crude material was subjected to column chromatography to afford tert-butyl N-[(3-chlorophenyl)[(isoquinolin-4-yl)carbamoyl]methyl]carbamate (3.0 g, 7.28 mmol, 32.7% yield).

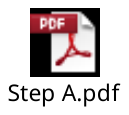

**Step B:**

CC(C)(C)OC(=O)NC(C(=O)NC=1C=NC=C2C=CC=CC12)C=3C=CC=C(Cl)C3.CC(=O)Cl>>NC(C(=O)NC=1C=NC=C2C=CC=CC12)C=3C=CC=C(Cl)C3

tert-Butyl N-[(3-chlorophenyl)[(isoquinolin-4-yl)carbamoyl]methyl]carbamate (3.0 g, 7.29 mmol) was dissolved in MeOH and acetyl chloride (1.72 g, 21.86 mmol, 1.56 ml, 3.0 equiv) was added at r.t. and heated at 50 ºC overnight. Then, the mixture was diluted with saturated K_2_CO_3_ solution, extracted with CHCl_3_. Organic layer was dried under sodium sulphate and evaporated to give 2-amino-2-(3-chlorophenyl)-N-(isoquinolin-4-yl)acetamide (2.1 g, 6.74 mmol, 92.4% yield).

**Step C:**

NC(C(=O)NC=1C=NC=C2C=CC=CC12)C=3C=CC=C(Cl)C3.OC(=O)[1C@H]1CC[1C@H](O1)C(=O)N2CCCC2.(CN(C)C(=[N+](C)C)N1N=[N+]([O-])C=2N=CC=CC12.F[P-](F)(F)(F)(F)F).CCN(C(C)C)C(C)C>>ClC=1C=CC=C(C1)C(NC(=O)[1C@@H]2CC[1C@@H](O2)C(=O)N3CCCC3)C(=O)NC=4C=NC=C5C=CC=CC45

rac-(2R,5S)-5-(Pyrrolidine-1-carbonyl)oxolane-2-carboxylic acid (621.63 mg, 2.92 mmol), ethylbis(propan-2-yl)amine (1.88 g, 14.58 mmol, 2.54 ml, 5.0 equiv) and 2-amino-2-(3-chlorophenyl)-N-(isoquinolin-4-yl)acetamide (1.0 g, 3.21 mmol) in DMF were cooled to 0 °C and hexafluoro-lambda5-phosphanuide 1-[(dimethylamino)(dimethyliminiumyl)methyl]-1H-[1,2,3]triazolo[4,5-b]pyridin-3-ium-3-olate (1.33 g, 3.5 mmol) was added. The reaction mixture was stirred at room temperature overnight. The reaction mixture was subjected to HPLC purification to afford rac-2-(3-chlorophenyl)-N-(isoquinolin-4-yl)-2-[(2R,5S)-5-(pyrrolidine-1-carbonyl)oxolan-2-yl]formamidoacetamide (148.39 mg, 292.69 µmol, 10% yield). The synthesis was repeated in large scale.

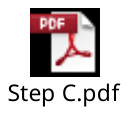

**Step D: Chiral resolution:**

Racemic rac-2-(3-chlorophenyl)-N-(isoquinolin-4-yl)-2-[(2R,5S)-5-(pyrrolidine-1-carbonyl)oxolan-2-yl]formamidoacetamide (500.00 mg) was subjected to chiral HPLC Column: Chiralpak AS-H (250 * 20 mm, 5 mkm); Mobile phase: Hexane-IPA-MeOH, 50-25-25. Flow Rate: 12 mL/min; Column Temperature: 24 ºC; Wavelength: 205 nm. RetTime (isomer A) = 9.4 min; RetTime (isomer B) = 21.11 min to afford 211.8 mg of rel-(2S,5R)-N-((R)-1-(3-chlorophenyl)-2-(isoquinolin-4-ylamino)-2-oxoethyl)-5-(pyrrolidine-1-carbonyl)tetrahydrofuran-2-carboxamide (Z5056159151, H3319657).

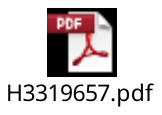

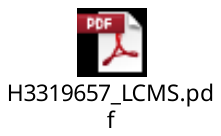

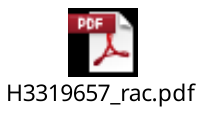

**The Synthesis of rel-(4R)-6-chloro-4-({[(3-fluoro-4-sulfamoylphenyl)methyl]amino}methyl)-N-(isoquinolin-4-yl)-3,4-dihydro-2H-1-benzopyran-4-carboxamide (Z5118588394, H3241102)**

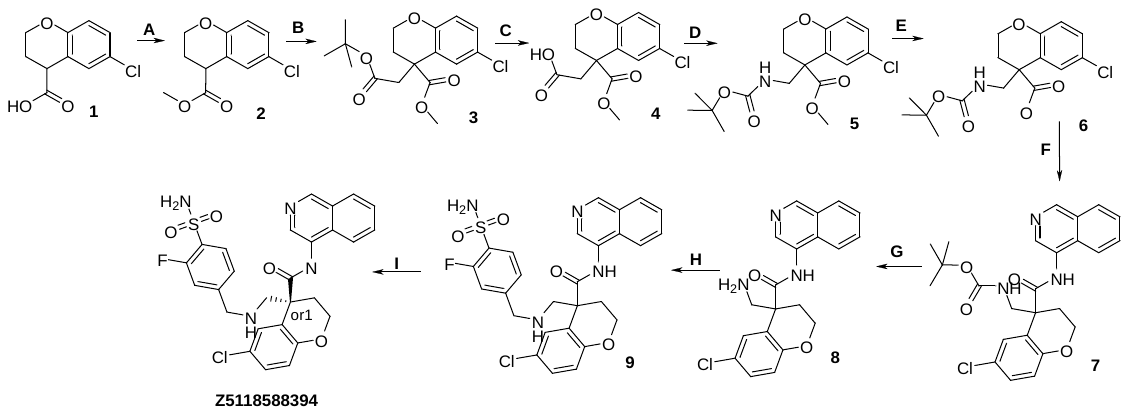

**Step A:**

6-Chloro-3,4-dihydro-2H-1-benzopyran-4-carboxylic acid (25.0 g, 117.58 mmol, l, 1.0 equiv) , trimethoxymethane (112.29 g, 1.06 mol) and 4-methylbenzene-1-sulfonic acid hydrate (223.65 mg, 1.18 mmol) were mixed in MeOH (250ml) and RM was heated 65°C overnight. Then solvent was evaporated and crude was diluted with MTBE and washed twice with saturated NaHCO3. Organic lawyer was separated, dried over Na2SO4 and solvent was evaporated under reduced pressure to give methyl 6-chloro-3,4-dihydro-2H-1-benzopyran-4-carboxylate (27.0 g, 95.0% purity, 113.17 mmol, 96.3% yield) as orange oil which was used directly in the next step without purification.

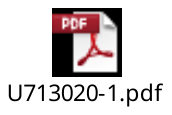

**Step B:**

Diisopropylamine (20.49 g, 202.54 mmol, 28.66 ml, 1.7 equiv) was dissolved in anhydrous THF (150ml) under Ar and cooled to -20°C. N-butyllithium (11.45 g, 178.71 mmol, 73.19 ml, 1.5 equiv) (28% solution in hexane) was added dropwise at this temperature and RM was allowed to warm to 0°C during 30 min. Then a solution of methyl 6-chloro-3,4-dihydro-2H-1-benzopyran-4-carboxylate (27.0 g, 119.14 mmol, l, 1.0 equiv) in THF (100ml) was added dropvise at 0°C and RM was allowed to warm to rt and stirred at rt for 2h. Then RM was cooled to -50°C and tert-butyl 2-bromoacetate (25.56 g, 131.05 mmol, 19.22 ml, 1.1 equiv) was added at this temperature. RM was stirred overnight at rt. Then, saturated NH4Cl solution was added and solvent was evaporated under reduced pressure. Crude was diluted with EtOAc and washed with water, saturated NH4Cl solution and brine. Organic lawyer was separated, dried over Na2SO4 and solvent was evaporated under reduced pressure to give crude which was dissolved in MTBE and filtered through a pad of silica and evaporated to give methyl 4-[2-(tert-butoxy)-2-oxoethyl]-6-chloro-3,4-dihydro-2H-1-benzopyran-4-carboxylate (37.0 g, 90.0% purity, 97.71 mmol, 82% yield) which was used directly in the next step without purification.

**Step C:**

Methyl 4-[2-(tert-butoxy)-2-oxoethyl]-6-chloro-3,4-dihydro-2H-1-benzopyran-4-carboxylate (37.0 g, 108.57 mmol) was dissolved in DCM (400ml) and 2,2,2-trifluoroacetic acid (86.66 g, 759.98 mmol, 58.67 ml, 7.0 equiv) was added. RM was stirred overnight. HNMR showed full conversion. Solvent was evaporated to obtain crude oil. This oil was reevaporated with toluene 3-5 times until it turned to solid. This solid was dried under oil pump to obtain 2-[6-chloro-4-(methoxycarbonyl)-3,4-dihydro-2H-1-benzopyran-4-yl]acetic acid (27.0 g, 94.84 mmol, 87.4% yield) which was used directly in the next step without purification.

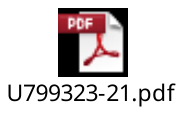

**Step D:**

2-[6-Chloro-4-(methoxycarbonyl)-3,4-dihydro-2H-1-benzopyran-4-yl]acetic acid (27.41 g, 96.27 mmol) , triethylamine (10.72 g, 105.9 mmol, 14.76 ml, 1.1 equiv) and anhydrous MgSO4 was suspended in dry toluene (500ml) under Ar and [azido(phenoxy)phosphoryl]oxybenzene (25.17 g, 91.46 mmol) was added dropvise. RM was heated 50°C for 1 h and then 90°C for 2h. Then absolute 2-methylpropan-2-ol (14.27 g, 192.54 mmol) was added and RM was refluxed overnight. HNMR showed full conversion. RM was cooled, diluted with EtOAc and washed with saturated NaHCO3 5 times. Organic lawyer was separated, dried over Na2SO4 and solvent was evaporated under reduced pressure to give crude product, which was purified by column chromatography (Interchim; 450g SiO2, Hex-MTBE% 0-10-100, flow rate = 46 mL/min. Rv=3-6,2 column equilibration Hexane. Incoming matter dissolved in chcl3) to obtain pure methyl 4-([(tert-butoxy)carbonyl]aminomethyl)-6-chloro-3,4-dihydro-2H-1-benzopyran-4-carboxylate (20.0 g, 95.0% purity, 53.4 mmol, 55.5% yield)

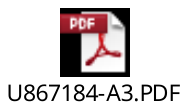

**Step E:**

Methyl 4-([(tert-butoxy)carbonyl]aminomethyl)-6-chloro-3,4-dihydro-2H-1-benzopyran-4-carboxylate (20.0 g, 56.21 mmol) was dissolved in MeOH (200ml) and water solution of lithium hydroxide monohydrate (7.08 g, 168.63 mmol) was added. RM was stirred overnight. LCMS showed full conversion. Solvent was evaporated and crude was diluted with water. Water phase was washed twice with MTBE and carefully acidified with NaHSO4 solution. Product was extracted 4 times with EtOAc. Organic lawyer was separated, dried over Na2SO4 and solvent was evaporated under reduced pressure to give 4-([(tert-butoxy)carbonyl]aminomethyl)-6-chloro-3,4-dihydro-2H-1-benzopyran-4-carboxylic acid (18.0 g, 95.0% purity, 50.03 mmol, 89% yield) which was used directly in the next step without purification.

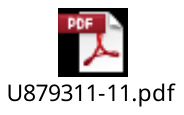

**Step F:**

A stirred mixture of 4-([(tert-butoxy)carbonyl]aminomethyl)-6-chloro-3,4-dihydro-2H-1-benzopyran-4-carboxylic acid (18.0 g, 52.66 mmol, l, 1.0 equiv) , isoquinolin-4-amine (8.73 g, 60.56 mmol) and 1-methyl-1H-imidazole (17.29 g, 210.63 mmol, 16.79 ml, 4.0 equiv) in dry ACN (300ml) was cooled in ice bath under Ar and Chloro-N,N,N,N-tetramethylformamidinium hexafluorophosphate (19.21 g, 68.46 mmol) was added. RM was stirred overnight. After complection of the reaction (by LCMS) solvent was evaporated and crude was diluted with EtOAc. Organic lawyer was washed with wated, saturated NaHCO3 solution, brine, dried over Na2SO4 and solvent was evaporated under reduced pressure to give crude product which was purified by column chromatography (CHCl3:ACN 80/20. 950g SiO2) to obtain tert-butyl N-(6-chloro-4-[(isoquinolin-4-yl)carbamoyl]-3,4-dihydro-2H-1-benzopyran-4-ylmethyl)carbamate (15.0 g, 95.0% purity, 30.45 mmol, 57.8% yield) .

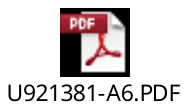

**Step G:**

Tert-butyl N-(6-chloro-4-[(isoquinolin-4-yl)carbamoyl]-3,4-dihydro-2H-1-benzopyran-4-ylmethyl)carbamate (14.77 g, 31.57 mmol) was dissolved in DCM (100ml) and hydrogen chloride (6.08 g, 166.69 mmol, 60.78 ml, 6.0 equiv) (10% solution in dioxane) was added. After complection of the reaction (monitored by LCMS) solvent was evaporated and crude was reevaporated with dioxane. Crude was diluted with diethyl ether and product was filtered as a pale-orange powder to obtain 4-(aminomethyl)-6-chloro-N-(isoquinolin-4-yl)-3,4-dihydro-2H-1-benzopyran-4-carboxamide dihydrochloride (12.0 g, 96.0% purity, 26.14 mmol, 94.1% yield)

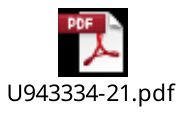

**Step H:**

4-(Aminomethyl)-6-chloro-N-(isoquinolin-4-yl)-3,4-dihydro-2H-1-benzopyran-4-carboxamide dihydrochloride (300.0 mg, 680.66 µmol) , 2-fluoro-4-formylbenzene-1-sulfonamide (145.85 mg, 717.81 µmol) and triethylamine (242.12 mg, 2.39 mmol) were mixed in anhydrous ACN and stirred for 1h. Then sodium cyanoboranuide (112.77 mg, 1.79 mmol) was added and mixture was stirred at rt overnight. 40% conversion was observed by LCMS. Saturated K2CO3 solution was added to RM and stirred for 30 min. Solvent was evaporated, crude was purified by reverse phase HPLC (2 runs) to give 6-chloro-4-([(3-fluoro-4-sulfamoylphenyl)methyl]aminomethyl)-N-(isoquinolin-4-yl)-3,4-dihydro-2H-1-benzopyran-4-carboxamide (33.0 mg, 85.0% purity, 50.54 µmol, 8.4% yield) .

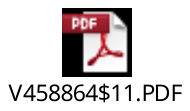

**Step I: Chiral separation**

Racemic 6-chloro-4-([(3-fluoro-4-sulfamoylphenyl)methyl]aminomethyl)-N-(isoquinolin-4-yl)-3,4-dihydro-2H-1-benzopyran-4-carboxamide (47.0 mg, 84.68 µmol) was separated by reverse phase chiral HPLC to obtain both enantiomers (isomer A 8mg; isomer B 20mg). Column: Chiralpak IC (250 * 20 mm, 5 mkm); Mobile phase : IPA-MeOH, 50-50. Flow Rate: 12 mL/min; Column Temperature: 24'C; Wavelength: 205 nm. RetTime (isomer A) = 11.59 min; RetTime (isomer B) = 15.39 min.

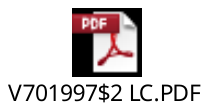

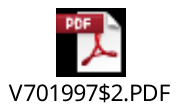

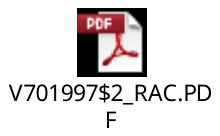

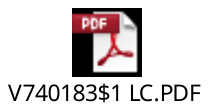

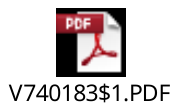

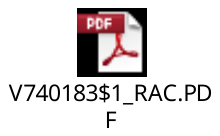

**The Synthesis of 6-chloro-4-[({[4-fluoro-3-(methylsulfamoyl)phenyl]methyl}amino)methyl]-N-(isoquinolin-4-yl)-3,4-dihydro-2H-1-benzopyran-4-carboxamide (Z4984330290)**

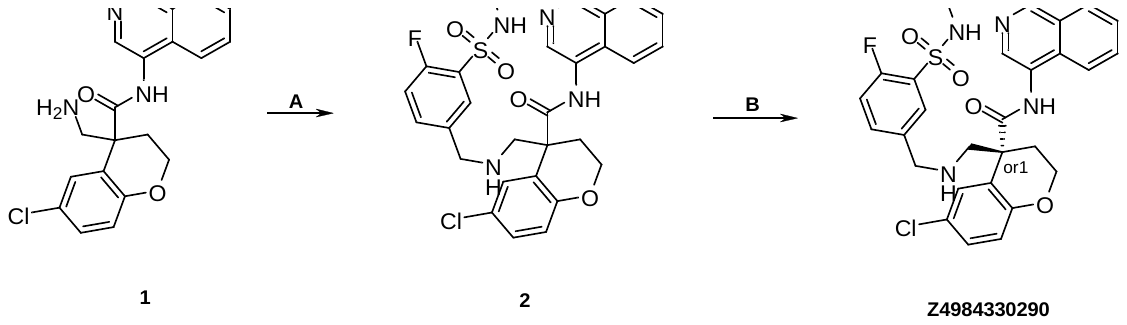

**Step A:**

4-(Aminomethyl)-6-chloro-N-(isoquinolin-4-yl)-3,4-dihydro-2H-1-benzopyran-4-carboxamide dihydrochloride (216.1 mg, 490.29 µmol) , 2-fluoro-5-formyl-N-methylbenzene-1-sulfonamide (160.0 mg, 736.59 µmol) and triethylamine (148.84 mg, 1.47 mmol, 210.0 µl, 3.0 equiv) were mixed in anhydrous ACN and stirred for 1h. Then sodium cyanoboranuide (154.06 mg, 2.45 mmol) was added and mixture was stirred at rt overnight. 57% conversion was observed by LCMS. Saturated K2CO3 solution was added to RM and stirred for 30 min. Solvent was evaporated, crude was purified by reverse phase HPLC (2 runs) to give 6-chloro-4-[([4-fluoro-3-(methylsulfamoyl)phenyl]methylamino)methyl]-N-(isoquinolin-4-yl)-3,4-dihydro-2H-1-benzopyran-4-carboxamide (39.0 mg, 90.0% purity, 61.68 µmol, 12.6% yield).

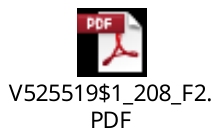

**Step B: Chiral separation**

Racemic 6-chloro-4-[({[4-fluoro-3-(methylsulfamoyl)phenyl]methyl}amino)methyl]-N-(isoquinolin-4-yl)-3,4-dihydro-2H-1-benzopyran-4-carboxamide (47.0 mg, 84.68 µmol) was separated by reverse phase chiral HPLC to obtain both enantiomers (isomer A 8mg; isomer B 20mg). Column: ChiralART YMC(250 * 20 mm , 5mkm); Mobile phase: Hexane-IPA-MeOH 50-25-25 Flow Rate: 12mL/min. RetTime (isomer A) = 11.01 min; RetTime (isomer B) = 16.29 min.

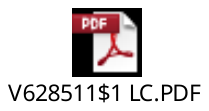

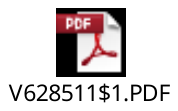

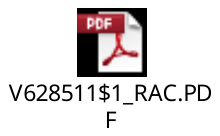

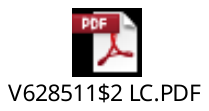

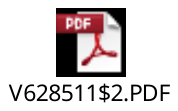

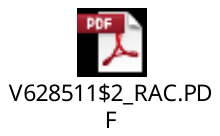
