## Supplementary Materials - Supplementary Figures for "Turning high-throughput structural biology into predictive inhibitor design"

### High throughput crystallography in predictive ligand design - Supplementary Information

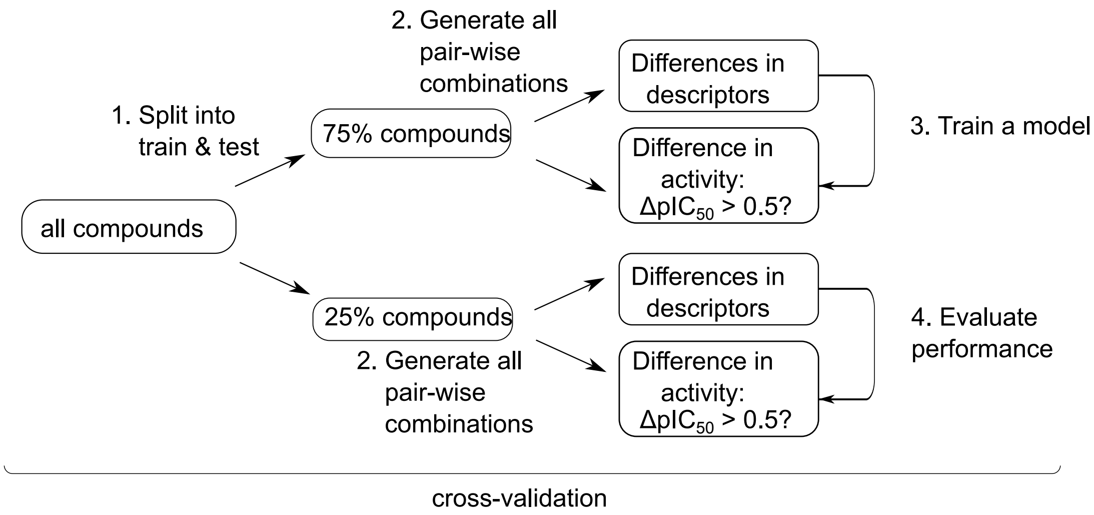


Figure S1. Overview of the model development process. The data was split into a training and a test set in 3:1 ratio (1) and pair-wise differences between the fingerprint and the activity value for all the compounds estimated (2). A model was trained that learned as a function of the differential fingerprint if two compounds differed by more than a half $pIC_{50}$ unit. The performance of the models was estimated on the test set. 10-fold cross-validation was used throughout the study unless otherwise specified.

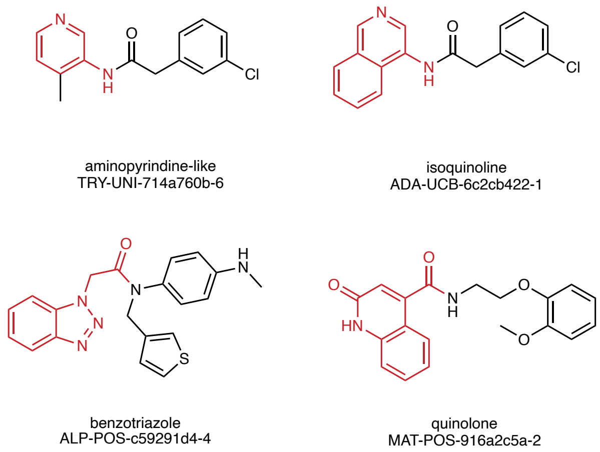


Figure 2: Figure S2. Division of the molecules into chemical series. Representative example from each of the four chemical series, with the salient chemical motif highlighed in red.

| AUROC values | Aminopyridine-like (n = 123) | Isoquinoline (n = 44) | Benzotriazole (n = 19) | Quinolone (n = 15) |
|---|---|---|---|---|
| Ligand-based learning | 0.64 ± 0.05 | 0.46 ± 0.07 | 0.42 ± 0.11 | 0.56 ± 0.10 |
| Docking-based learning | 0.51 ± 0.09 | 0.62 ± 0.02 | 0.79 ± 0.02 | 0.82 ± 0.03 |
| Docking w/ structure-based learning | 0.80 ± 0.01 | 0.71 ± 0.01 | 0.80 ± 0.01 | 0.78 ± 0.02 |

Figure S3. The performances of the three models for each of the four scaffolds. The values correspond to the AUROC scores on the left-our training data by 20 models trained with bootstrapping.

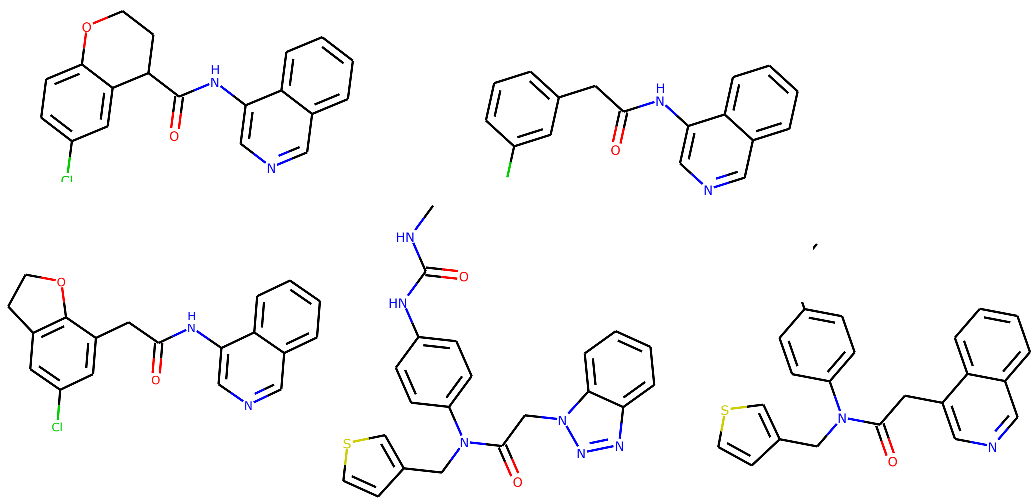

Figure S4. Structures of the five highly potent non-covalent compounds from the COVID moonshot campaign that were used as the reference when evaluating the relative rankings of each of the compounds in the virtual library.
